# Cerebrovascular super-resolution 4D Flow MRI – using deep learning to non-invasively quantify velocity, flow, and relative pressure

**DOI:** 10.1101/2021.08.25.457611

**Authors:** E. Ferdian, D. Marlevi, J. Schollenberger, M. Aristova, E.R. Edelman, S. Schnell, C.A. Figueroa, D.A. Nordsletten, A.A. Young

## Abstract

The development of cerebrovascular disease is tightly coupled to changes in cerebrovascular hemodynamics, with altered flow and relative pressure indicative of the onset, development, and acute manifestation of pathology. Image-based monitoring of cerebrovascular hemodynamics is, however, complicated by the narrow and tortuous vasculature, where accurate output directly depends on sufficient spatial resolution. To address this, we present a method combining dedicated deep learning and state-of-the-art 4D Flow MRI to generate super-resolution full-field images with coupled quantification of relative pressure using a physics-driven image processing approach. The method is trained and validated in a patient-specific *in-silico* cohort, showing good accuracy in estimating velocity (relative error: 12.0 ± 0.1%, mean absolute error (MAE): 0.07 ± 0.06 m/s at peak velocity), flow (relative error: 6.6 ± 4.7%, root mean square error (RMSE): 0.5 ± 0.1 mL/s at peak flow), and with maintained recovery of relative pressure through the circle of Willis (relative error: 11.0 ± 7.3%, RMSE: 0.3 ± 0.2 mmHg). Furthermore, the method is applied to an *in-vivo* volunteer cohort, effectively generating data at <0.5mm resolution and showing potential in reducing low-resolution bias in relative pressure estimation. Our approach presents a promising method to non-invasively quantify cerebrovascular hemodynamics, applicable to dedicated clinical cohorts in the future.

## I. INTRODUCTION

Changes in regional hemodynamics are intimately coupled to the manifestation of cerebrovascular disease, making the quantification of flow and pressure critically important for improving diagnostics. Variations in pressure throughout the cerebrovasculature have been particularly highlighted in a number of clinical scenarios: the functional impact of intracranial atherosclerosis linked to regional changes in intravascular pressure [1], the likelihood of cerebral aneurysm growth related to regional pressure gradients [2], and experimental work showing altered pressure variations in arteriovenous malformations [3]. While transcranial Doppler or 2D phase-contrast magnetic resonance imaging (PC-MRI) provide limited information on regional flow, it is through time-resolved three-dimensional phase-contrast magnetic resonance imaging (4D Flow MRI) that full-field hemodynamic mapping can be achieved [4]. 4D Flow MRI has been used in a number of studies to capture cerebrovascular flow phenomena [5], and in combination with physics-informed image processing, quantification of relative pressure is permitted [6]. However, spatial resolution is insufficient to accurately quantify both cerebrovascular flow [7] and relative pressures [6], where vessel diameters ≤3 voxels, or dx ≥0.75 mm in the circle of Willis have been shown to result in significant biases while current clinical systems are limited to around dx = 0.5-1 mm. Similarly, because image noise scales with resolution, high-resolution acquisitions require extended scan times, making them clinically cumbersome. In summary, there remains a definite need for effective approaches to achieve higher-resolution flow imaging for cerebrovascular hemodynamic assessment.

To address the need for improved spatial resolution, high-Tesla approaches have been proposed [8, 9], but are inherently limited to specialized imaging systems. Image-guided computational fluid dynamics (CFD) modelling has also been explored [10, 11], however, this approach generally puts high demand on available computational resources, and further depends on boundary conditions typically requiring additional specialized imaging protocols [11].

As an alternative to these deterministic approaches, deep learning methods have recently been applied in the field of medical image enhancement. For MRI, deep learning methods have been proven to enable data denoising [12], artefact compensation [13], and to generate super-resolution anatomical reconstructions of the brain [14]. For flow-based MRI, 2D studies have shown the ability to generate accelerated reconstructions of phase-contrast images [15], as well as enable automatic flow quantification over network-segmented flow domains [16]. For 4D Flow MRI, Ferdian et al. [17] proposed the so-called 4DFlowNet to generate super-resolution 4D Flow MRI data from low-resolution input, with the network trained on synthetic pairs of low/high-resolution images generated from aortic CFD simulations. Other alternatives include Rutkowski et al. [12] using a convolutional neural network (CNN), and Fathi et al. [18] using a Physics-Informed Neural Network (PINN), both generating super-resolution 4D Flow images using CFD input data as ground truth for training. Whilst 4DFlowNet was only tested on large-vessel aortic flows, both the CNN and the PINN-based alternatives were implemented on either phantom-data resembling cerebrovascular flow, or on selected *in-vivo* sets. However, no extended quantitative analysis has been performed for *in-vivo* usage in a cerebrovascular setting. Furthermore, neither of the above-mentioned networks (2D or 3D) have been tested with respect to functional pressure measurements, and it remains unknown whether pressure changes through the image domain are maintained or even improved by applying any of these super-resolution procedures.

The aim of this study is therefore to assess whether a dedicated cerebrovascular super-resolution network could improve estimates of regional cerebrovascular velocities and flows, and in particular, whether functional relative pressure estimates could be improved by means of super-resolution image conversion. To achieve this, the existing super-resolution network 4DFlowNet is re-purposed to the cerebrovascular space using dedicated sets of multiresolution input training data, originating from patient-specific CFD models with relevant image features (magnitude, noise level) extracted from conjunctive, clinically acquired 4D Flow MRI. After validating recovery of velocity, flow, and functional relative pressures *in-silico*, the re-purposed network is applied to an *in-vivo* cohort of subjects scanned at multiple resolutions, assessing the potential of super-resolution imaging in a more clinical setting. In summary, our study explores the potential of non-invasive super-resolution imaging for cerebrovascular usage, providing improved estimates of clinically relevant functional hemodynamics metrics throughout the cerebrovasculature.

## II. METHODS

### A. Deep learning framework for cerebrovascular super-resolution flow imaging

#### 1) Deep learning network architecture

To achieve super-resolution flow images, we utilize the deep residual network structure of 4DFlowNet [17]; a previously published network validated for large-vessel aortic flows. Briefly, the architecture is based on a central upsampling layer (using bilinear interpolation) surrounded by a series of stacked residual blocks (RB), with preceding RBs denoising and pre-processing the input, and subsequent RBs refining and sharpening the predicted output. As input, both low resolution magnitude and velocity phase image patches were utilized. As output, super-resolution velocity patches were generated.

We used a similar design for the original 4DFlowNet structure [17], with the following specific changes introduced for its application on cerebrovascular flow data:

1. Patch input size was changed from an original 16-voxel cube, to a 12-voxel cube, accounting for the smaller vessel sizes encountered in the cerebrovascular space.
2. The original hyperbolic tangent activation functions at the output layers were switched to linear activation functions. This was introduced to aid the network in reducing overfitting whilst still allowing for unbounded output values.
3. The gradient terms were removed from the loss function, following improvements observed in near-wall velocity estimates in preliminary data assessment.

The modified network was trained using an Adam optimizer, with a learning rate set to 10-4. Batch sizes of 20 were used for training, with training completed after 60 epochs. The network was implemented using Tensorflow 2.0 [19], utilizing a Keras backend.

#### 2) Loss function definition

For the loss function, the optimization target was set to minimize the mean squared error (MSE) between the generated super-resolution images, and the paired high-resolution input data. The voxel-wise loss was defined as the mean of the squared sum of differences between Cartesian velocity components, (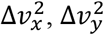 and 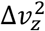), given as

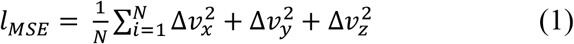

where N is the total number of voxels in the assessed image domain. To compensate for imbalances between fluid and static tissue regions within a singular patch, the MSE was calculated separately for fluid and static tissue in each region.

Lastly, to avoid network overfitting, an L2 regularization term was included. The complete loss function was given as

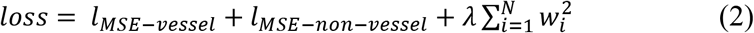

where *l*_*MSE−vessel*_ and *l*_*MSE−non−vessel*_ are the voxel-wise MSE loss in fluid and static tissue, respectively, and λ is a set coefficient (equal to 5 · 10^−7^) regularizing the network weights *w*_*i*_.

#### 3) Cerebrovascular training and testing data

To train the super-resolution network, sets of low and high resolution flow images needed to be collected. Whilst acquired, matched, integer pairs of clinical 4D Flow MRI data would represent a theoretically ideal training set, in practice it is very difficult to obtain such high-resolution, high-SNR, artefact-free *in-vivo* ground truth data suitable for training. Instead, we here propose a separate set of synthetic 4D Flow MRI originating from patient-specific cerebrovascular flow simulations. To improve clinical relevance, simulated data are combined with reference *in-vivo* scans, from which realistic noise levels and relevant reference magnitude images can be extracted.

##### a) Patient-specific *in-silico* data

As a basis for training, anatomically accurate patient-specific CFD models of the arterial cerebrovasculature were used, providing both realistic velocity, flow, and reference pressure fields data [11].

In short, models were created using a combination of time-of-flight (TOF) MRI, 2D phase contrast (PC) MRI, and MRI arterial spin labelling (ASL) [20], covering the vasculature from the aortic root to the circle of Willis (CoW). A pulsatile velocity profile derived from PC-MRI was prescribed at the inlet of the aortic root. Each outlet was coupled to a 3-element Windkessel lumped parameter model and calibrated using a combination of PC-MRI and ASL perfusion data [11]. 3D models were meshed using tetrahedral elements, and the incompressible Navier-Stokes equations solved iteratively using a stabilized finite-element formulation. Nodal velocity and pressure data were extracted after periodicity was reached (≥4 cardiac cycles). The modelling and analysis were performed using the validated open-source framework CRIMSON [21]. Further details on setup and model validation can be found in [11]. Data from four different image sets were generated:

<u>Subject 1</u> presenting without evidence of cerebrovascular disease, although exhibiting an incomplete CoW through right and left posterior communicating artery hyperplasia.

<u>Subject 2</u> presenting with a severe stenosis in the right proximal internal carotid artery (ICA, 70-99% based on velocity criteria from duplex ultrasound) and a complete CoW.

<u>Subject 3a</u> presenting with a bilateral carotid stenosis (80-90% in the right proximal ICA, and 60% in the left proximal ICA, based on CTA image criteria), and a CoW exhibiting right P1 segment and distal right vertebral artery hypoplasia.

<u>Subject 3b</u> being the same subject as 3a following surgical re-opening of the stenosis at the right proximal ICA.

From the above, synthetic 4D Flow MRI data was generated by sampling the nodal CFD output onto a uniform voxelized image grid. With the aim of covering varying spatial scales, data were generated for spatial samplings of dx = 1.5, 1.0, 0.75, 0.5, and 0.375 mm isotropic, respectively (allowing for high/low resolution pairs of 1.5/0.75; 1.0/0.5; and 0.75/0.375 mm). A time step of dt = 1 ms was used to increase the amount of input data for training. Data was consistently extracted for a region-of-interest (ROI) centered around the intracranial vessels. An illustration of one of the utilized models is shown in Figure 1.

**Figure 1.**
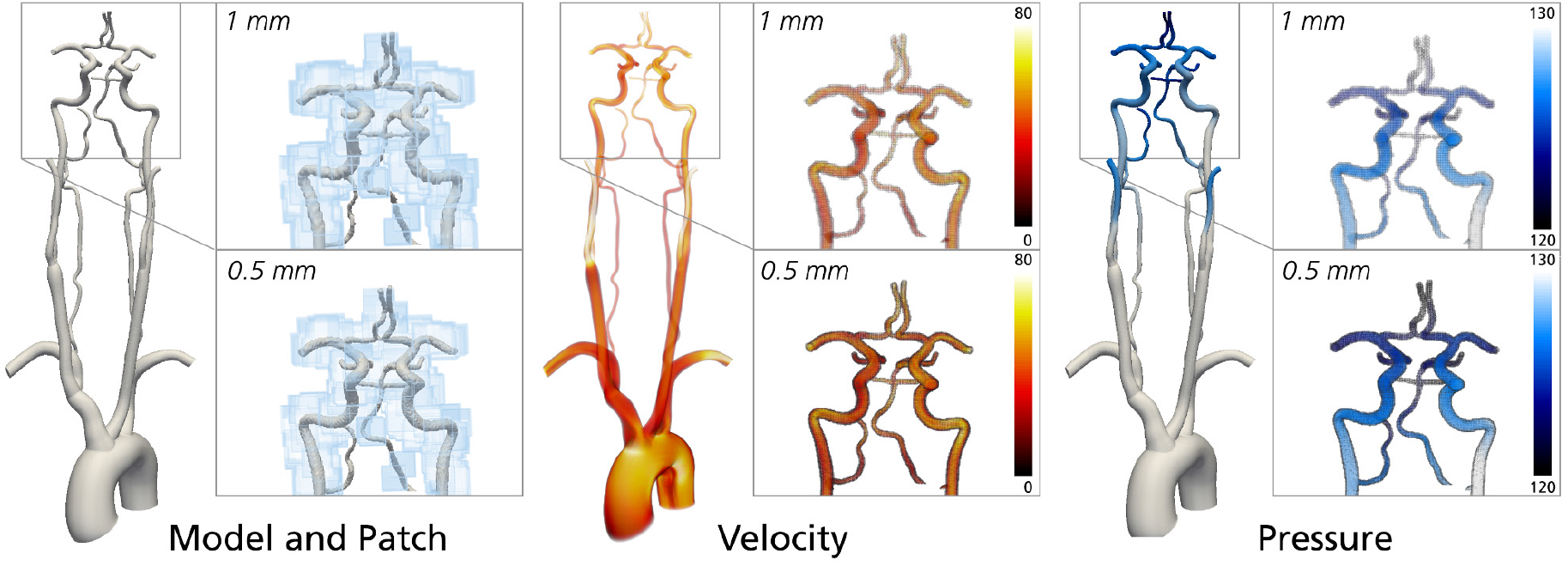
Overview of the *in-silico* input used for re-training of the 4DFlowNet network, showing one of the four used models (Subject 3b). From left to right: model overview and patch generation through the proximal cerebrovascular ROI; velocity field (color range 0 - 80 cm/s); pressure field (color range 120 - 130 mmHg). Note that examples are shown for the low/high-resolution pair of 1.0/0.5 mm isotropic.

##### b) Cerebrovascular *in-vivo* data

Using a cohort of 8 healthy volunteers (2 women, 6 men, 55 ± 18 years), MRI acquisitions were performed at 3T (Siemens Magnetom Skyra, Erlangen, Germany) using a 20-channel head/neck coil. Centering a ROI around the CoW, acquisitions started with a TOF MRA sequence (TR = 21 ms; TE = 3.6 ms; flip angle = 18°), followed by 4D Flow MRI (prospective k-t GRAPPA dual-venc (130/45 cm/s) acquisitions [22], dt = 95-104 ms). Flow images were acquired at two different resolutions: dx = 1.1 mm isotropic, and dx = 0.8 mm isotropic. Scan times were 10-15 minutes for all sequences, respectively. In all instances, data were corrected for concomitant gradient fields, eddy currents, and noise. All clinical acquisitions followed institutional review board (IRB) approval and informed consent.

##### c) Training and testing data patch generation

To enhance clinical relevance of the training data, synthetic 4D Flow MRI from a) were transformed into clinical-quality equivalents. In short, realistic velocity-to-noise ratios (VNR) were extracted from the clinical data in b), equaling approximately VNR = 5.67 ± 1.64 at dx = 1.1 mm, and VNR = 2.97 ± 0.78 at dx = 0.8 mm. With data from a) treated as effective phase information, and with clinical magnitude images from b) used as reference, clinical-level noise was added to the synthetic 4D Flow MRI through k-space downsampling, extracting complex numbers from the synthetic phase and clinical magnitude images, respectively. Note that such noise was added to the low-resolution dataset only, resulting in a network tasked not only with increasing resolution, but also removing noise.

To generate a larger number of training sets from the limited (n = 4) number of models, the FOV was split into patches of restricted spatial extent. Specifically, from each temporal frame patches of 123 voxels were extracted from random positions within the FOV (enforcing a minimum flow region of >5%). Visualization of the distribution of patches is shown in Figure 1. For every patch, data augmentation by rigid cartesian rotations (90/180/270°) were applied.

Data from Subjects 1 and 2 were selected for training with a total of 42,900 patches, Subject 3a for validation consisting of 2,730 patches, and Subject 3b for testing. Training was performed on a Titan X GPU with 12GB memory. With training performed for 60 epochs, lasting approximately 30 minutes each, complete training took about 30 hours. Super-resolved velocity fields were predicted on a patch-basis, with complete volumes reconstructed by stitching patches together with a stride of n = 8 voxels in each Cartesian direction, with n being an arbitrary patch size configurable during inference. Note that 4 voxels were stripped from each patch side, reducing data to the patch center

### B. Validation of super-resolution performance, and recovery of cerebrovascular relative pressure

#### 1) *In-silico* validation of super-resolution velocity, flow, and relative pressure

To validate performance of the super-resolution network, the *in-silico* models and corresponding synthetic 4D Flow MRI data from Section II.A.3 (a) were utilized. Performance was evaluated with respect to both super-resolved velocity fields and derived flows, as well as functional recovery of relative pressures using coupled physics-informed image processing.

##### a) Validation of super-resolution velocity and flow

For the super-resolved velocity fields, linear regression analysis was performed against reference high-resolution velocity data from the CFD analysis, assessing Cartesian velocity components and velocity magnitudes separately. Bland-Altman plots of the same data were also extracted to assess potential network bias. For general quantification, assessment of mean absolute error (MEA), root mean square error (RMSE), cosine similarity, absolute magnitude error, and relative magnitude error were all performed, with the latter extracted as per

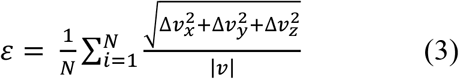

with 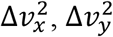, and 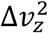 being Cartesian velocity components.

Furthermore, flow rates through three different planes cutting through sections of the right ICA, mid-ICA, and MCA were also compared between super-resolved and high-resolution reference synthetic 4D Flow MRI data. Quantification of RMSE and relative errorwere also performed against high-resolution reference flow from the CFD analysis.

##### b) Validation of super-resolution relative pressure

A key component of our study was to assess whether network-based super-resolution images also enabled accurate extraction of conjunctive, functional relative pressures. A variety of methods exist to derive relative pressures from image velocity data, each with specific method assumptions and applicability in the cerebrovascular space. Here we use the virtual work-energy relative pressure (vWERP) method, which allows for arbitrary probing through narrow and bifurcating structures [23], with catheter-based validation underlining the method’s potential. vWERP has also been applied in a cerebrovascular setting, indicating promising abilities whilst highlighting the importance of sufficient spatial resolution [6].

With details provided in previous work [23], vWERP originates from a virtual work-energy form of the Navier-Stokes equations, derived by introducing an auxiliary virtual field w, and evaluating the resulting expression over the fluid domain of interest, Ω. Doing so, relative pressures can be derived as:

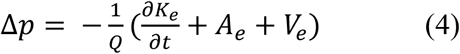

with

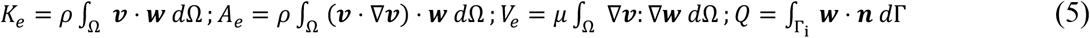

Here, each term represents different *virtual* energy components, including virtual kinetic energy (*K*_*e*_), virtual advective energy rate (*A*_*e*_), virtual viscous energy dissipation (*V*_*e*_), and the virtual flow (*Q*) going through a selected inlet plane (Γ_i_). Introducing ***w*** as a divergence-free field with ***w*** = 0 at all domain wall boundaries, relative pressures can then be extracted directly from the imaged flow field ***v***.

Using *v*WERP, relative pressures were estimated over four different cerebrovascular sections in each synthetic 4D Flow MRI dataset, respectively: *<u>left / right ICA</u>*, going from the cranial end of the cervical ICA to the mid-section of the petrous ICA, and *<u>left / right ICA-middle cerebral artery (MCA)</u>*, going from the mid-section of the petrous ICA to midway along the M1-segment of the MCA. Based on previous analysis [6], estimations were performed on low/high resolution pairs of 1.0/0.5 and 0.75/0.375 mm, as well as on corresponding super-resolution data. In all instances, data were extracted with temporal sampling of dt = 40 ms, to approximate a clinically realistic acquisition.

Just as in Section II.B.1(a), linear regression analysis was performed for super-resolved relative pressures against reference high-resolution pressure field data originating from the simulated CFD output. Bland-Altman plots were also extracted to assess potential estimation bias. For general quantification, assessment of RMSE, cosine similarity, and relative error was also performed, as per Section II.B.1(a).

#### 2) *In-vivo* implementation and possibilities for clinical cerebrovascular super-resolution

Adding to the validation in Section II.B.1, super-resolved velocity fields were also generated and assessed in the clinical 4D Flow MRI data from Section II.A.3(b). Super-resolution upsampling was performed by a factor of two on all datasets (converting 1.1 to 0.55 mm, and 0.8 to 0.4 mm, respectively).

##### a) Estimation of super-resolution velocity and flow

Native and super-resolved flow fields were qualitatively compared to assess visual correspondence. Although data were not acquired in integer resolution pairs, through-plane flow rates at the proximal section of the left and right MCAs were still compared between resolution sets to quantify differences between native and super-resolved resolutions, as well as changes in velocity-to-noise ratio (VNR).

##### b) Estimation of super-resolution relative pressure

To assess relative pressures in the *in-vivo* data, similar ICA-MCA sections as the ones used in the *in-silico* analysis were identified. To achieve this, vessel segmentation was first performed using a previously published analysis framework [24]. Second, inlet and outlet planes for the relative pressure estimations were positioned based on relevant anatomical landmarks along the right and left ICA and MCA, with planes visually co-aligned between resolutions (1.1 and 0.8 mm, respectively). With planes and segmentations created, vWERP was used to extract relative pressures in all subjects. Whilst lacking reference pressures, extracted measures were compared over different resolutions, assessing linear correlations and Bland-Altman plots between the different sets (with and without super-resolution, respectively).

## III. RESULTS

### A. *In-silico* validation of super-resolution 4D Flow MRI

#### 1) Validation of super-resolution velocity and flow

Complete evaluation was performed on one test subject (Subject 3b), using 1 mm input data (low resolution, LR) to generate super-resolution equivalents at 0.5 mm (SR), comparing output quality against high-resolution (HR) reference data at the same 0.5 mm resolution. As apparent in Figure 2, significant noise reduction is achieved in the SR velocity fields. Furthermore, SR flow rates indicate slight overestimation at the proximal-most (A) section (mean shift of -0.33 ± 0.14 mL/s), whilst showing a similar but opposite underestimation of flow in the more distal (B) and (C) sections (0.34 ± 0.16 mL/s, and 0.33 ± 0.12 mL/s, respectively). Relative differences are however kept <10.3 % over the evaluated sections (Figure 2 and Table 1). Isolating peak flow rates in all models, slight error reduction is seen for conversion from LR (RMSE = 0.74 mL/s, relative error = 9.0 ± 6.2%) to SR (RMSE = 0.56 mL/s, relative error = 6.6 ± 4.7%).

**Figure 2.**
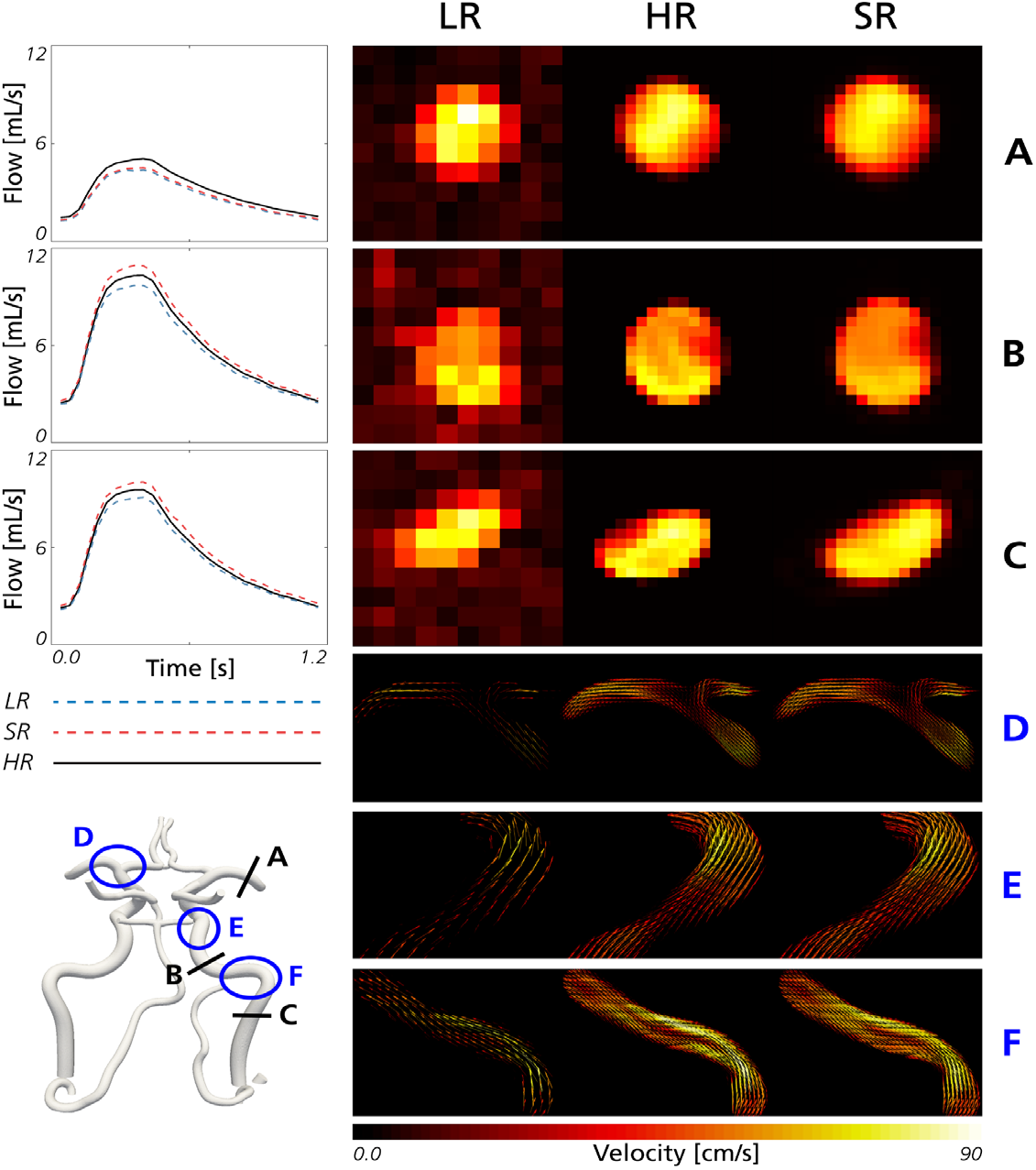
Comparison between low resolution (LR), high resolution (HR), and super resolution (SR) images at three different intersecting planes (A-C) and three different regional sections (D-F) all through the ICA-MCA. Insets are showing the selected regions in magnified form and with views rotated to highlight velocity vectors. Comparison of flow rates through the intersecting planes (A-C) are also shown. Note that the model insert at the bottom left is shown dorsally.

**Table 1.**
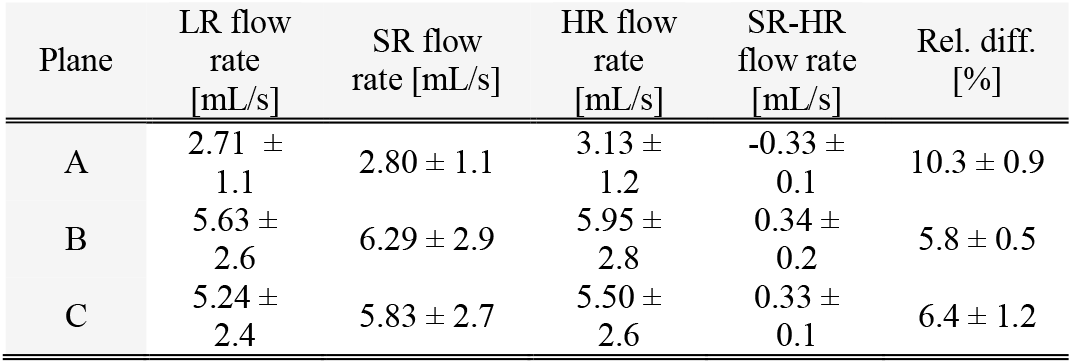
Flow rate measurements on Subject 3b for the right MCA, mid-ICA, and ICA. For all sections, results were measured by averaging 3 parallel cross-sectional slices.

Figure 3 shows linear regression plots and Bland-Altman representations for generated super-resolution velocities. In general, excellent correlations are observed between SR and HR velocities, with linear regression slopes and correlation coefficients of k>0.91 and R2>0.96 reported for the vessel core region (all voxels apart from the outermost fluid layer), and k>0.90 and R2>0.76 for the vessel wall region (the outermost layer of fluid voxels). Slightly lower values are seen for velocity magnitudes (k=0.82 and R2=0.79 for core; k=0.69 and R2=0.52 for wall), although the Bland-Altman output corroborates the quality of the results, with minimal bias indicated (consistent deviations of <0.02 m/s).

**Figure 3.**
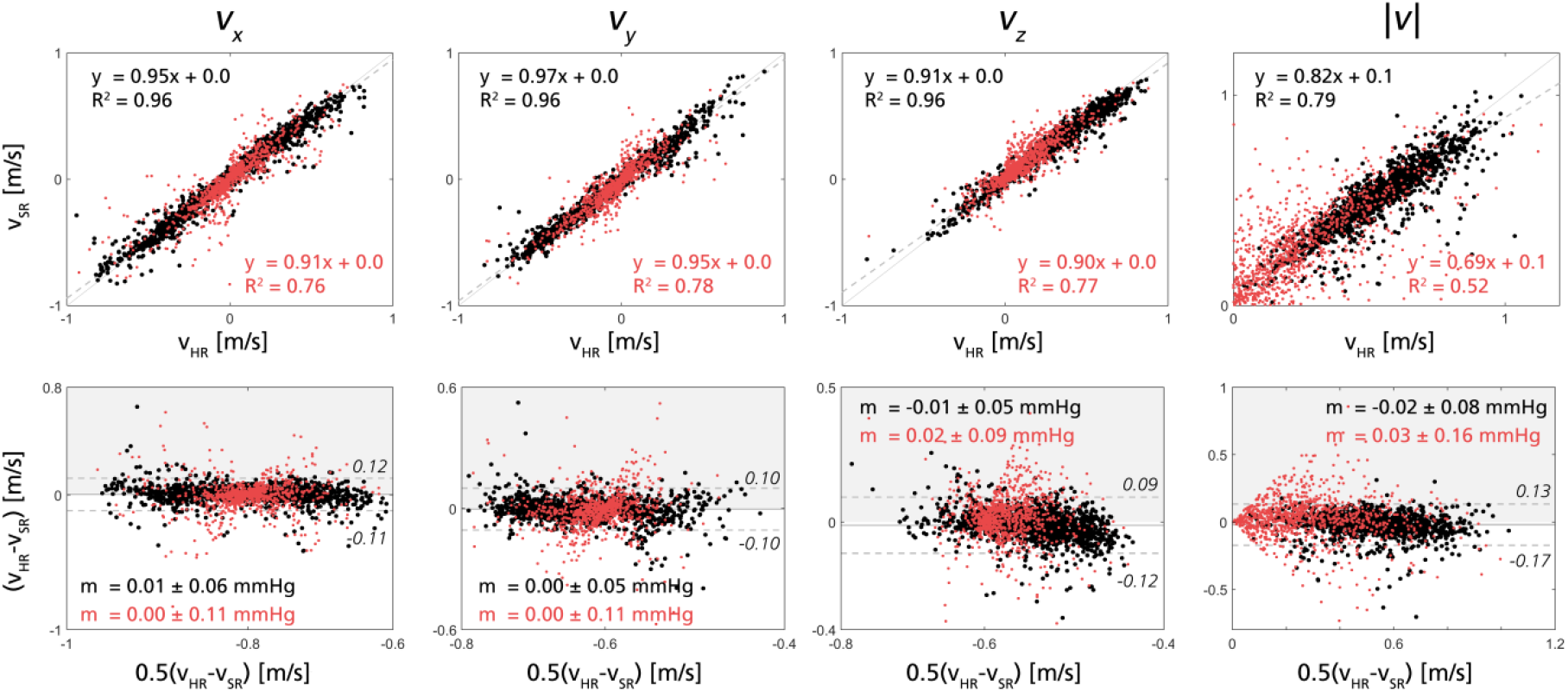
Top: Regression plot for each of the velocity components (vx, vy, and vz) and velocity magnitude between ground truth and super-resolved image during the peak flow for *in-silico* test case (Subject 3b). Bottom: Bland-Altman plot for each of the velocity components during peak flow. The plots show 5% of the data points (randomly selected) within the vessel core (black) and vessel wall (red), respectively

Isolating peak velocity magnitudes, measures in both vessel core (MAE = 0.07 ± 0.06 m/s, relative error = 12.0 ± 0.07%, cosine similarity = 0.99 ± 0.06) and vessel wall regions (absolute error = 0.12 ± 0.11 m/s, and cosine similarity = 0.95 ± 0.11) confirm the trends noted above. Similar numbers are also observed for 0.75/0.375 mm resolution sets, as shown in Supplementary Material A.

#### 2) Validation of super-resolution relative pressure

Figure 4 shows linear regression and Bland-Altman plots for estimations of relative pressure across different resolutions and all models (example relative pressure traces are also given in Supplementary Material B). Overall, significant underestimation is observed at LR (1 mm), whilst accurate estimates are reported at the HR (0.5 mm) setting. Importantly, distinct improvements in functional relative pressures are observed for the super-resolved SR fields as compared to the LR input: relative error in peak relative pressure decreasing from 23.3 ± 14.9 % at LR, to 11.0 ± 7.3 % at SR, with 5.1 ± 2.3 % at reference HR. Similarly, the RMSE for the entire time series goes from 1.1 ± 1.7 mmHg at LR, to 0.3 ± 0.2 mmHg at SR, compared to 0.2 ± 0.1 mmHg at HR. Quantitative output for an ICA-MCA sections across all different models are given in Table 2.

**Figure 4.**
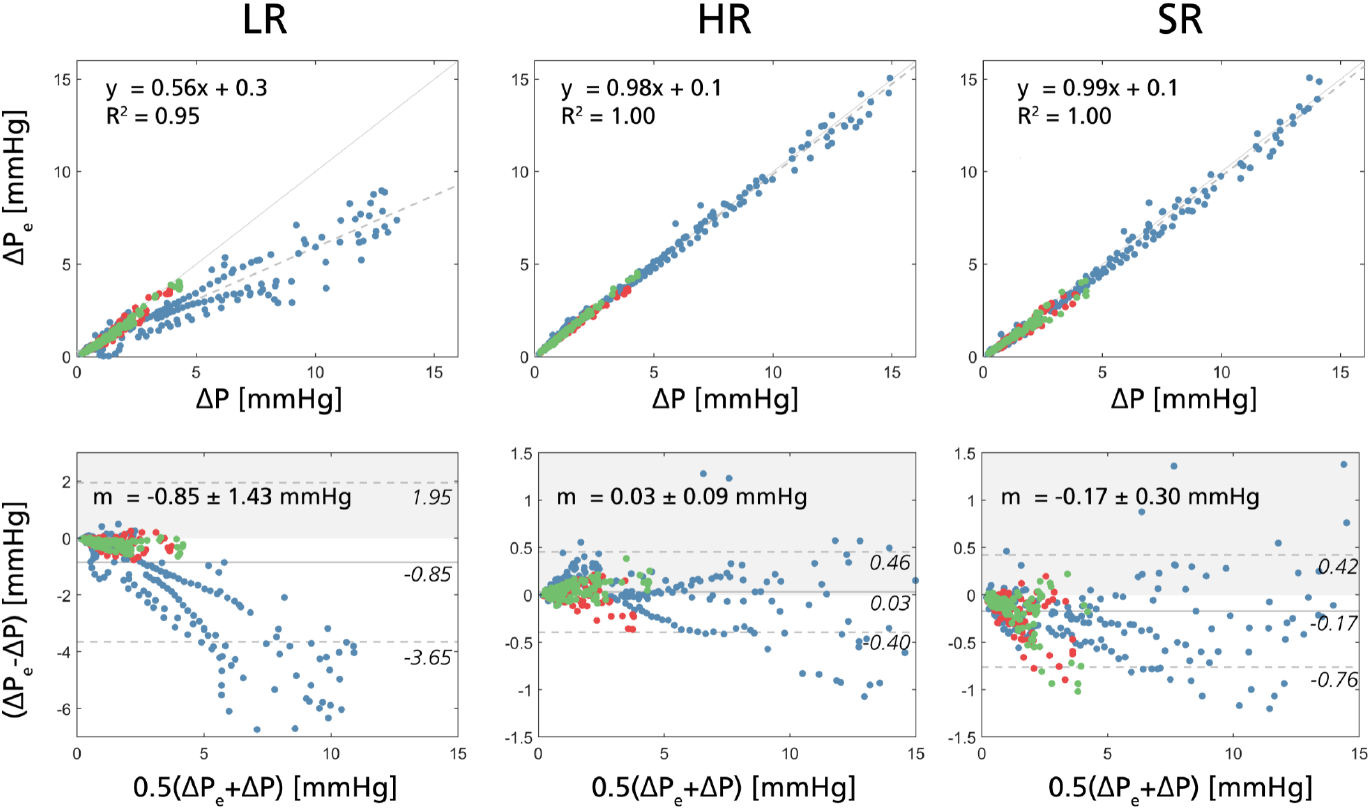
Linear regression (top row) and Bland-Altman plots (bottom row), comparing relative pressure estimates to reference CFD equivalents using low resolution data (LR, 1 mm, left column), high resolution data (HR, 0.5 mm, middle column), and super-resolution data (SR, converting 1 mm to 0.5 mm, right column). The colors depict different data sets (training in blue (Subject 1 and 2), validation in red (Subject 3a), testing in green (Subject 3b)).

**Table 2.** Image-based peak relative pressure measurements through the right ICA-MCA section four all different subjects.

| Model | LR peak<br>$\Delta p$<br>[mmHg] | SR peak<br>$\Delta p$<br>[mmHg] | HR peak<br>$\Delta p$<br>[mmHg] | SR-HR<br>peak $\Delta p$<br>[mmHg] | Rel. diff.<br>[%] |
| --- | --- | --- | --- | --- | --- |
| 1 | 7.39 | 13.93 | 13.11 | 14.04 | 0.82 |
| 2 | 6.78 | 12.97 | 12.51 | 13.00 | 0.46 |
| 3a | 2.51 | 2.91 | 2.81 | 2.88 | 0.10 |
| 3b | 2.35 | 2.80 | 2.66 | 2.71 | 0.14 |

The above is also confirmed in Figure 4 with conversion from LR to SR increasing the linear regression slope from k = 0.56 to 0.99, representing a virtual 1:1 correlation to ground truth relative pressures (k = 0.98 at HR for reference). Likewise, the mean bias shift in the LR set (mean shift of -0.85 ± 1.43 mmHg) is significantly reduced by conversion into SR data (mean shift of -0.17 ± 0.30 mmHg). The HR data show no estimation bias (mean shift of 0.03 ± 0.22 mmHg). Notice that similar improvements are observed when converting 0.75 mm base resolution sets into super-resolution equivalents (at 0.375 mm), with complete data for this analysis shown in Supplementary Material A.

#### 3) Comparison between original and re-trained networks

Super-resolution predictions were also computed using the original aortic 4DFlowNet. With detailed results provided in Supplementary Material C, the aortic network shows deviations from ground truth HR data when it comes to super-resolved velocity components (linear regression slopes and correlation coefficients of k>0.73 and R2>0.55, and k>0.51 and R2>0.29 are reported for vessel core and wall regions, respectively). The original aortic 4DFlowNet also shows lower accuracy for the recovery of cerebrovascular relative pressures (k = 0.87 against reference data, and a mean bias shift of -0.41 ± 0.58 mmHg). Peak relative pressure estimates are given at a relative error of 14.8 ± 11.9 %, and a RMSE of 0.5 ± 0.6 mmHg – all consistently higher than what is reported for the repurposed cerebrovascular 4DFlowNet. Again, complete data are provided in Supplementary Material C.

### B. *In-vivo* implementation of cerebrovascular super-resolution 4D Flow MRI

#### 1) Estimation of super-resolution velocity and flow

For the *in-vivo* dataset, visual inspection confirmed qualitative improvement with regards to noise reduction and data appearance of the generated super-resolved 4D Flow MRI data (see Figure 5). Specifically, VNR showed a 4-times increase in the 0.55 mm SR data (going from VNR = 5.67 ± 1.64 at dx = 1.1 mm to VNR 24.20 ± 11.28 at dx = 0.55 mm), and a 3-times increase in the 0.4 mm SR data (going from VNR = 2.97 ± 0.78 at dx = 0.8 mm to VNR 9.29 ± 4.25 at dx = 0.4 mm).

**Figure 5.**
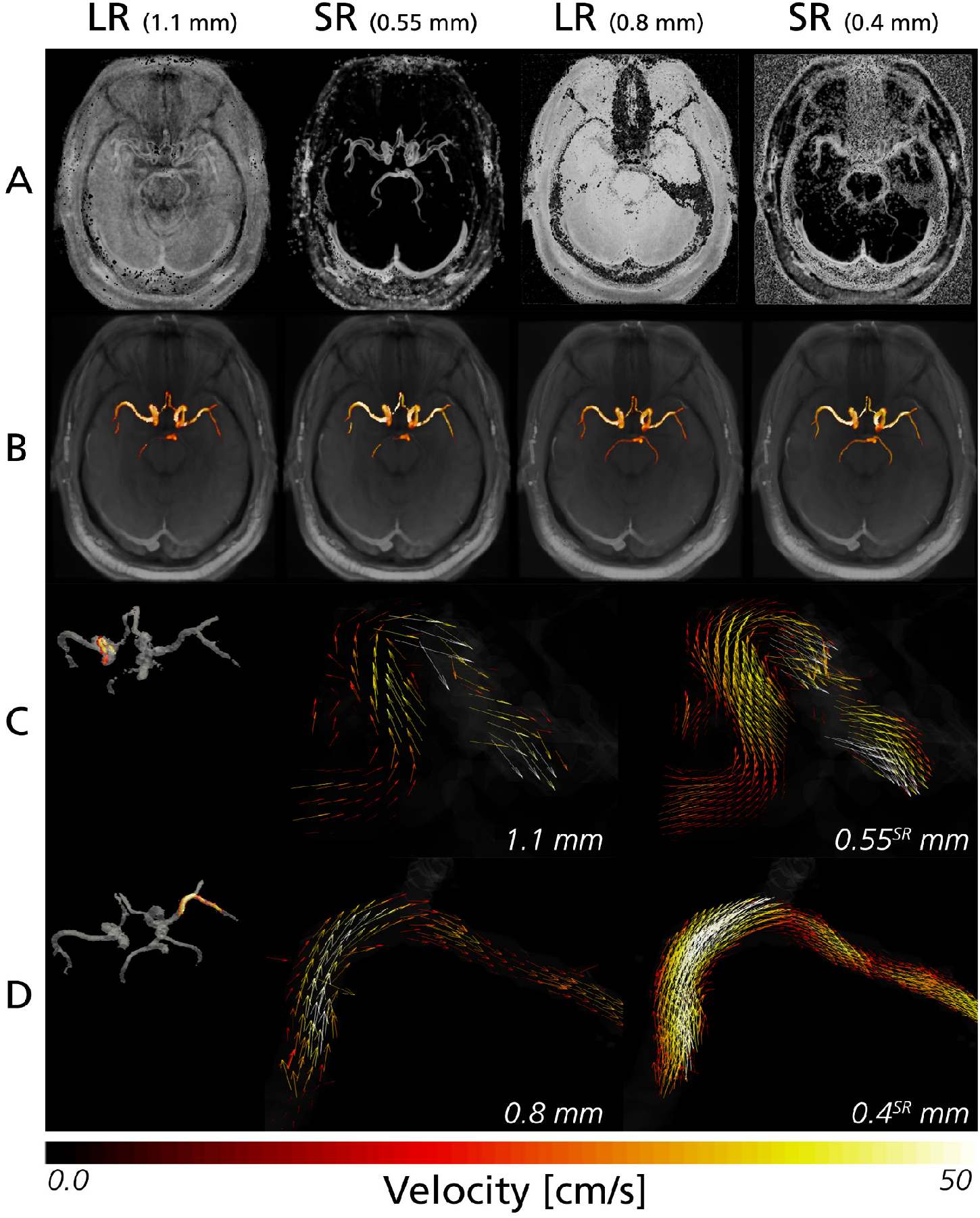
Visual comparison of an *in-vivo* case at low (LR) and super-resolution (SR) given for both sets of dx=1.1 and 0.55, and 0.8 and 0.4 mm, respectively. Improvements in VNR are apparent in the super-resolved phase images (A) as well as in the flow visualizations (B). Direct velocity vectors comparison are given for a section through the right MCA for the paired low resolution (1.1 mm)/super-resolution (0.55 mm) in (C), and for the paired low resolution (0.8 mm)/super-resolution (0.4 mm) in (D), with vectors shown projected onto a visual2D plane. In general, broad view of the velocity vectors only reveal minor differences between resolution sets, although detailed view reveals velocity vectors conforming more to the anatomy of the vessel in the super-resolved images, including at the near-wall regions

Assessing flow rates through the left and right MCAs, the clinical base resolution data indicated a flow rate range of 0.65 to 7.13 mL/s and peak flow rates of 4.96 ± 1.52 mL/s at dx = 1.1 mm, compared to a slightly reduced range of 0.67 to 5.53 mL/s and peak flow rates of 3.47 ± 1.01 mL/s at dx = 0.8 mm. Converting to SR equivalents (dx = 0.55 mm and 0.4 mm, respectively) flow rates are only modestly modified, with slight downregulation observed in both datasets (flow range of 0.58 to 6.93 mL/s and peak flow rates of 4.39 ± 1.56 mL/s at dx = 0.55 mm; flow range of 0.64 to 5.13 mL/s and peak flow rates of 3.32 ± 0.91 mL/s at dx = 0.4 mm).

#### 2) Estimation of super-resolution relative pressure

Relative pressures were derived for all *in-vivo* subjects and sections. Overall, estimates were within the range of -0.6 to 6.0 mmHg for the 1.1 mm data, with peak relative pressures at 2.9 ± 1.6 mmHg, compared to a range of -0.1 to 6.8 mmHg for the 0.8 mm data, with peak relative pressures at 3.8 ± 1.8 mmHg. Converting to SR, the ranges changes with estimates getting closer to one another: SR data at dx = 0.55 mm (input at dx = 1.1 mm) exhibiting a range of -0.7 to 5.9 mmHg with peak relative pressures at 2.6 ± 1.4 mmHg; SR data at dx = 0.4 mm (input at dx = 0.8 mm) exhibits a range of -0.5 to 4.3 mmHg with peak relative pressures at 2.9 ± 1.1 mmHg.

Although lacking *in-vivo* reference pressure, Figure 6 shows linear regression and Bland-Altman plots comparing LR and HR data to its SR equivalents. At base resolutions (LR vs. HR) a systematic bias shift in relative pressure is observed between the two resolutions (k = 0.64; R2 = 0.81; mean shift = -0.93 ± 0.93 mmHg). Converting to super-resolved equivalents, however, the shift is reduced, although without completely recovering a 1:1 correlation between the two datasets (k = 0.81; R2 = 0.77; mean shift = -0.47 ± 0.72 mmHg).

**Figure 6.**
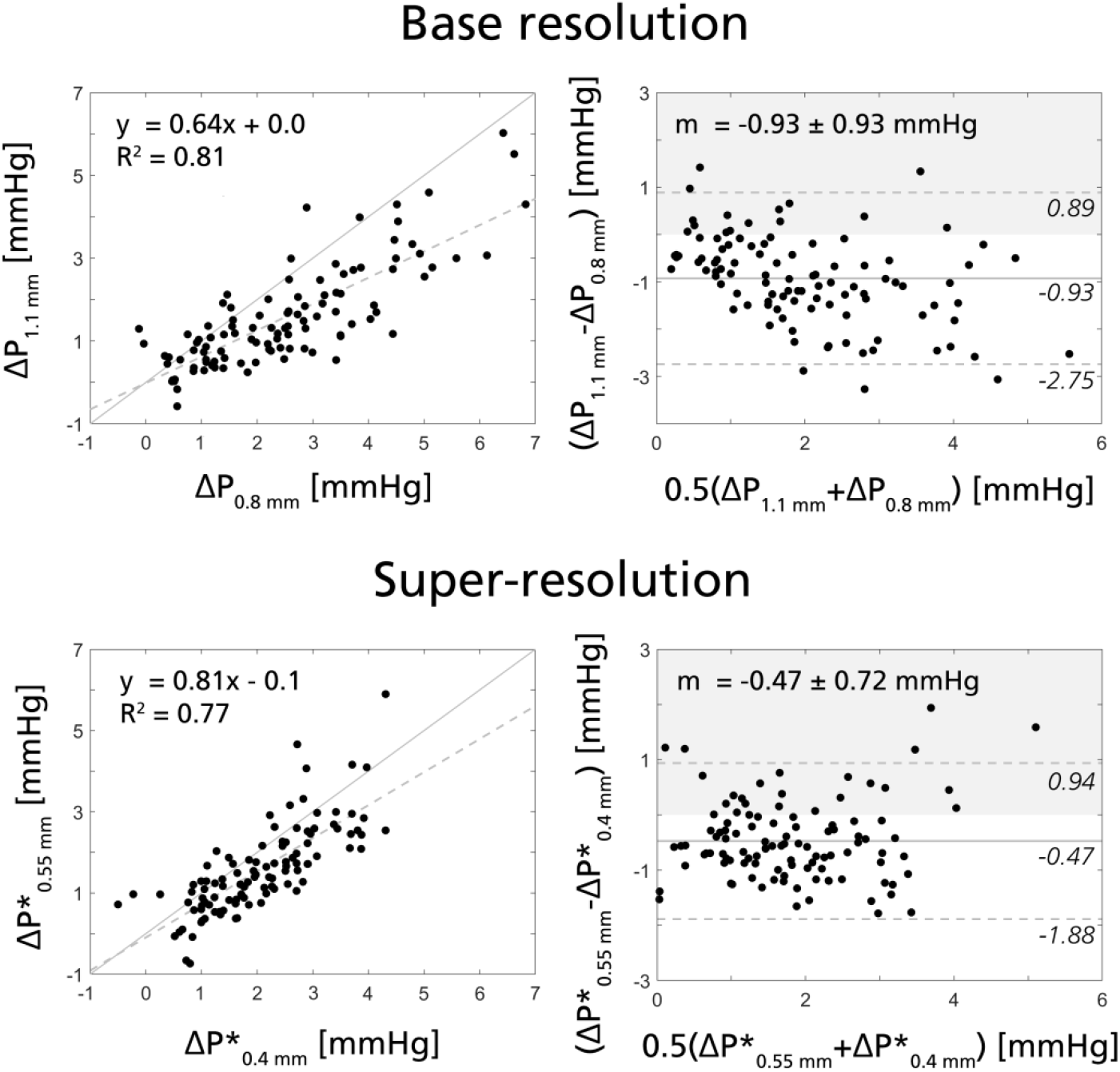
Linear regression and Bland-Altman plots for the *in-vivo* cerebrovascular 4D Flow MRI data, showing the relationship between relative pressure estimated at base resolutions (ΔP, two left-most plots, comparing 1.1 mm and 0.8 mm data) and at equivalent super-resolutions (ΔP*, two rightmost plots, comparing super-resolved 0.55 mm vs. 0.4 mm data).

## IV. DISCUSSION

In this study, we evaluated the utility of super-resolution 4D Flow MRI in the setting of cerebrovascular hemodynamics, showcasing how super-resolved intracranial velocity fields and regional flows can be recovered from low-resolution input data using a re-trained version of the 4DFlowNet architecture. Furthermore, we showed how super-resolution 4D Flow MRI in combination with the physics-informed vWERP algorithm successfully recovers functional relative pressures through regional cerebrovascular sections, with the super-resolved data effectively reducing estimation bias otherwise observed in the low-resolution input data. With non-invasive cerebrovascular assessment intrinsically complicated by the narrow and tortuous vasculature, our results highlight the potential of super-resolution 4D Flow MRI to improve quantitative functional cerebrovascular hemodynamic assessment.

### A. *In-silico* validation of cerebrovascular super-resolution 4D Flow MRI to quantify velocity, flow, and relative pressure

*In-silico* super-resolved flows and velocity fields both conform closely to high-resolution reference data. For super-resolved velocities, slightly reduced accuracy was identified along near-wall voxels. This behavior is similar to what has been previously reported [12], and is not entirely surprising: near-wall voxels suffer from reduced input information (being surrounded by ‘information-depleted’ static tissue), and will be inherently linked to reduced signal quality. Dedicated neural networks have been explored for the recovery of near-wall velocities in 2D flow data [25], although application in 4D Flow MRI data remains to be performed.

Furthermore, a major part of our work focused on whether super-resolved flow fields enable accurate estimation of functional relative pressures; an entity directly dependent on utilized spatial resolution [6]. As reported in Section III.A.2), the combination of a super-resolution network (4DFlowNet) and a physics-informed analysis algorithm (vWERP) allow for accurate estimation of cerebrovascular relative pressures. This not only indicates the utility of the vWERP algorithm but also highlights that the 4DFlowNet architecture allows for accurate estimation of the complete fluid mechanical environment, with precise recovery of both velocity and velocity gradients needed to accurately extract relative pressures.

Another benefit of the repurposed 4DFlowNet is the ability to significantly improve VNR. Deterministic multi-venc sequences have been explored to enhance VNR [22], however, using a post-processing super-resolution approach in principle enables maintained signal quality even at reduced scan times, as highlighted in other super-resolution work [12].

### B. Re-training 4DFlowNet for cerebrovascular usage

The importance of re-training is highlighted in Section III.A.3 and Supplementary Material C, where distinct performance differences are highlighted between the original (aortic) and repurposed (cerebrovascular) 4DFlowNet. Here, it is important to appreciate the fundamental differences in input training data that exist between the aortic and the cerebrovascular 4DFlowNet. In the original work, patches containing purely aortic flows from CFD were shown during training, with hemodynamics dominated by transient flows [26] guided through a large vessel structure. On the contrary, cerebrovascular hemodynamics is a joint resultant of transient, advective, and viscous behavior [6], with flow restricted by the narrow, tortuous vasculature. Additionally, the cerebrovascular training data contain synthetically generated magnitude images, as such carrying more realistic image properties and noise characteristics. Hence, the original network was never exposed to patches containing the same image characteristics, or entailing similarly small vessels or tortuous near-wall gradients, and performance is likely reduced when attempting cerebrovascular data recovery.

The fact that re-training resolved estimation bias also demonstrates that the core 4DFlowNet architecture is robust to different types of flows, and that it is rather the information contained in the training data (i.e. vessel sizes, noise characteristics) that determines final performance. This also indicates that further re-training might be necessary if attempting super-resolution imaging in yet another cardiovascular domain (e.g. intracardiac flow fields), although as long as anatomical structures are similar in size (e.g. cerebral vs. hepatic vessels) maintained accuracy is plausible. To overcome the need for constant retraining, one could envision combining training data from multiple domains to create a network handling both large and small vessel anatomies, as well as fast and slow flows. The performance of such a network, however, remains to be determined.

### C. *In-vivo* feasibility of cerebrovascular super-resolution 4D Flow MRI to quantify flow, velocity, and relative pressure

In Section III.B, super-resolution images and functional relative pressure estimations were performed in a select *in-vivo* cohort. Although ground truth high-resolution scans or reference pressure measurements were unavailable, the behavior indicated *in-silico* seems replicated *in-vivo*. Specifically, super-resolved flow fields did not introduce any bias shifts, and estimates of both flows and relative pressures indicate slight convergence at upsampled resolutions. Nevertheless, even though derived relative pressure magnitudes coincide with what has been reported in previous cerebrovascular work [27], a desired 1:1 relation between resolutions is not achieved. Here, comparably coarse temporal resolution (dt ≥ 95 ms), cardiovascular variations between scans, or temporal intra-scan mis-match could all contribute to this slight discrepancy. Further validation of *in-vivo* work would be beneficial to understand the clinical translation of the combined 4DFlowNet and vWERP approach.

### D. Contextualizing cerebrovascular super-resolution 4D Flow MRI

It is worth contrasting our re-purposed 4DFlowNet to previously published work within the same space. Whilst few studies exist attempting super-resolution recovery of directly imaged flow [12, 17], only a handful have attempted the same for functional hemodynamic recovery. Kissas et al. [28] proposed a PINN-based network to recover absolute pressure in simplified arterial model sections; however, application in cerebrovascular geometries was never attempted. Shit et al. [29] similarly proposed the PINN-based ‘Velocity-to-Pressure’ net; however, super-resolution abilities were never included. In comparison, our work combines the super-resolution utility of 4DFlowNet with the functional recovery of the physics-informed deterministic vWERP approach, being previously benchmarked across different cardiovascular domains, including the cerebrovasculature [6, 23, 30].

Continuing into the cerebrovascular space, a few very recent works have shown how merging physics-informed analysis, machine learning, and imaging can have particular promise for improving non-invasive cerebrovascular assessment. Fathi et al. [18] used a patient-specific PINN to recover regional flow and pressure from input 4D Flow MRI, promising virtually unrestricted spatiotemporal refinements on recovered velocity fields. Similarly, Rutkowski et al. [12] recently presented a CNN-based network to reconstruct super-resolution 4D Flow MRI in a cerebrovascular setting, using patient-specific in-vitro models for both training and testing. Along these very same lines, our work also highlights the significant potential of super-resolution 4D Flow MRI in the cerebrovascular space. Within this setting, our study extends these previous works by showing how a combination of super-resolution utilities with the physics-informed vWERP algorithm provides accurate recovery of relative pressures, overcoming inherent resolution bias otherwise observed in clinical-level image sets [6] and allowing for the accurate recovery of this established biomarker through the challenging cerebrovascular space. Whilst technical differences exist in utilized network design or loss function, the combination of our data and the above reviewed works all point to the increasing interest shown in network-driven super-resolution 4D Flow MRI, with the cerebrovascular space being a prime target of where such utilities can have direct clinical impact.

### E. Limitations

A number of limitations are worth pointing out. Firstly, clinical *in-vivo* validation against catheter based pressure data remains to be performed. Acquiring invasive pressure data in the cerebrovascular space is challenging as intracranial arterial catheterization still awaits regulatory approval in the US. Furthermore, clinical validation of super-resolution utilities is inherently limited in clinical practice. With both 4DFlowNet and the vWERP algorithm validated in other domains [23, 30], its potential in improving cerebrovascular quantification is evident. Still, experimental validation in patient-specific in-vitro models (as recently attempted in other super-resolution work [12]) or in a pre-clinical setting would bring important additional information as to the clinical utility of the presented work.

Secondly, a modest number of *in-silico* models were used for training, where additional data could enhance network versatility. Similarly, combining the original aortic and cerebrovascular datasets could generate a more general-purpose utility, although performance of such would have to be evaluated separately.

Thirdly, it is worth noting that the training of the super-resolution network also depends on the accuracy of the utilized CFD models to capture realistic cerebral flow and pressure. Realistic CFD modeling of cerebral flow is generally challenging due to difficulties in assigning patient-specific boundary conditions. In this work, however, we overcame these challenges by using a previously presented CFD calibration strategy based on cerebral perfusion (non-selective ALS) and flow (PC-MRI) data [11]. Specifically, the utilized CFD models were validated by comparing the blood supply in the CoW against territorial perfusion data from vessel-selective ALS, where observed high correlations underline the accuracy and applicability of the utilized models.

Lastly, a practical limitation is the increasing data storage required by the super-resolution conversion. Due to the uniformly sampled data representation, a two-fold resolution increase leads to an eight-fold increase in disk space usage. Adaptable grid representations or graph-based networks [31] may offer improved future possibilities circumventing this issue, and may be explored in future work.

### F. Clinical outlook and future work

The expansion of quantitative hemodynamic imaging for cerebrovascular applications promises improved clinical abilities [1-3], and the usage of super-resolution 4D Flow MRI presents an effective way of quantifying such hemodynamic markers in the brain, with our work highlighting its accurate recovery of both direct and functional hemodynamic metrics. Importantly, super-resolution imaging circumvents intrinsic obstacles otherwise related to non-invasive cerebrovascular flow quantification (limited spatial coverage; challenging vascular anatomies; etc.), and its clinical potential is therefore particularly evident within this vascular domain.

Numerous, future directions can be envisioned to extend and clarify the capabilities highlighted in our study: additional training data expanding network capabilities, modified architecture improving predictions in near-wall regions, or extended clinical validation against acquired 4D Flow MRI or experimentally derived invasive catheter data. Clinically oriented studies evaluating the potential of super-resolution imaging to improve clinical risk stratification by means of improved relative pressure estimations could also be envisioned in the future. Nevertheless, our data highlights the potential of super-resolution 4D Flow MRI and coupled physics-informed image analysis in the cerebrovascular space.

## V. CONCLUSION

In this study, we have shown how dedicated super-resolution 4D Flow MRI and physics-informed image analysis can together be effectively used to accurately quantify cerebrovascular hemodynamics, including regional velocities, flows, and functional relative pressures. Using dedicated patient-specific *in-silico* data, we have shown how the existing 4DFlowNet network can be effectively repurposed into the cerebrovascular space, successfully converting low-resolution input data into high-resolution equivalents with maintained precision and effective noise-reduction. Furthermore, in combination with the physics-informed deterministic image analysis algorithm vWERP, we have shown how conversion into super-resolution data successfully reduces estimation biases in functional relative pressures otherwise observed in the utilized low-resolution input data. Lastly, implementation in an exemplary *in-vivo* cohort shows how improvements in velocity-to-noise-ratio, preserved flow, and converging relative pressures estimates are achievable in a clinical setting.

