## Supplementary Material for "Cerebrovascular super-resolution 4D Flow MRI – using deep learning to non-invasively quantify velocity, flow, and relative pressure"

#### A. Validation of super-resolution performance over additional resolution sets

Corroborating the results in Section III.A.1 and III.A.2 similar analysis was performed over a second set of in-silico resolutions converting input data at dx = 0.75 mm to super-resolution data at dx = 0.375 mm (comparing against reference high-resolution data sampled directly from the equivalent CFD solution).

##### 1) Estimation of super-resolution velocity

As shown in Supplementary Figure 1 and in agreement with the results provided over the 1 mm/0.5 mm resolution pair in the main manuscript, excellent correlations are observed between super-resolution and high resolution data over all velocity components. Consistently, linear regression slopes and correlation coefficients are k>0.91 and R2>0.96 for the vessel core region, and k>0.95 and R2>0.74 for the vessel wall region. The Bland-Altman output also indicate no bias shifts introduced by the super-resolved data, with deviations of <0.05 m/s with limits of agreements <0.15 m/s across all components and regions. Isolating peak velocity magnitudes, measures in both vessel core (MAE = 0.07 ± 0.06 m/s, relative error = 14.38 ± 0.06%, cosine similarity 0.99 ± 0.06) and vessel wall regions (MAE = 0.12 ± 0.11 and cosine similarity 0.94 ± 0.11) similar with the 0.5/1.0 mm counterpart.

*First authorship is shared between E.F. and D.M., with both authors contributing equally to the work.
†Last authorship is shared between D.A.N and A.A.Y., with both authors contributing equally to the work.

E. F. holds a New Zealand Heart Foundation Scholarship, Grant No. 1786. D.M. holds a Knut and Alice Wallenberg Foundation scholarship for postdoctoral studies at Massachusetts Institute of Technology. J.S. is supported by a University of Michigan Rackham Predoctoral Fellowship. M.A. was supported by a Ruth L. Kirschstein National Research Service Award (NIH F30 HL140910) and the Northwestern – Medical Science Training Program (NIH T32 GM815229). E.R.E. was funded in part by NIH R01 49039. A.A.Y. acknowledges core funding from the Wellcome/EPSRC Centre for Medical Engineering (WT203148/Z/16/Z) and the London Medical Imaging and AI Centre for Value-Based Healthcare. D.N. would like to acknowledge funding from the Engineering and Physical Science Research Council (EP/N011554 and EP/R003866/1).

E.F. and A.A.Y. are with the University of Auckland, Auckland 1142 New Zealand. D.M. and E.R.E. are with the Massachusetts Institute of Technology, Cambridge, MA 02139 USA. J.S., C.A.F, and D.N. are with the University of Michigan, Ann Arbor, MI 48109, USA. M.A. and S.S. are with Northwestern University, Chicago, IL 60611, USA. S.S. is also with the University of Greifswald, Greifswad 17489, Germany. D.N. and A.A.Y. are also at King's College London, London, SE1 7EH, UK.

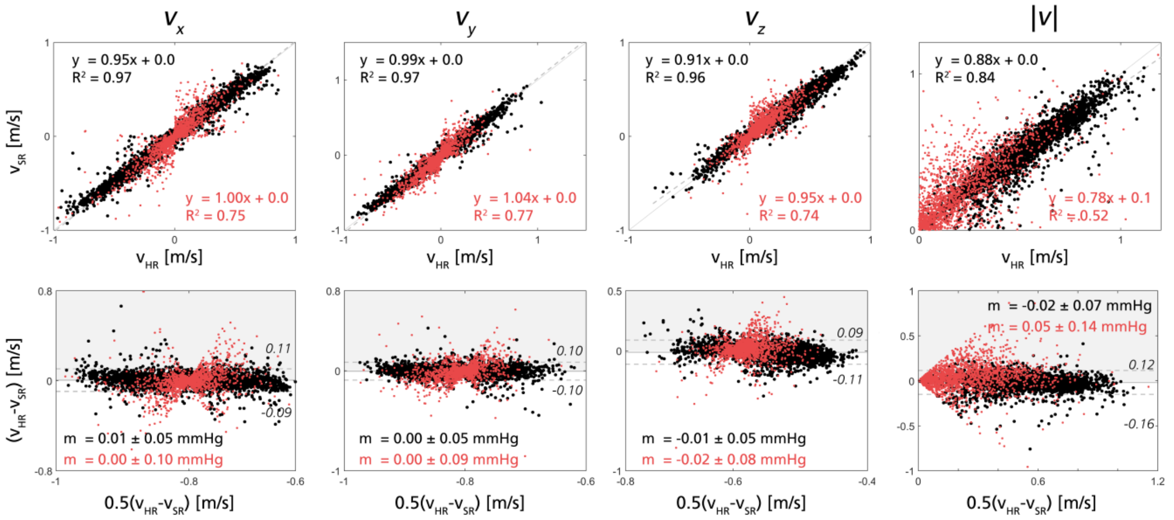


**Supplementary Figure 1** - Regression plot for each of the velocity components (Vx, Vy, and Vz) and velocity magnitude between ground truth and super-resolved image during the peak flow for in-silico test case (Subject 3b), using the additional resolution set of 0.75 mm (LR) and 0.375 mm (SR). Bottom: Bland-Altman plot for each of the velocity components during peak flow. The plots show 5% of the data points (randomly selected) within the vessel core (black) and vessel wall (red), respectively.

#### 2) Estimation of super-resolution relative pressure

Furthermore, Supplementary Figure 2 shows how conversion into super-resolution data mitigates underestimation in relative pressures, with a linear regression slope of changing from k = 0.83 at low resolution to k = 0.98 at super-resolution (to be compared with k = 1.02 at reference high-resolution). Bland-Altman assessments also supplements this same data, where the spread in estimates are reduced with super-resolution, albeit with a slightly remaining underestimation bias (mean shift of -0.25 ± 0.57 mmHg at low resolution; mean shift of -0.24 ± 0.24 mmHg at super-resolution; mean shift of 0.08 ± 0.17 mmHg at reference high resolution).

### B. Example super-resolution relative pressure traces

Complementing Section III.A.2, Supplementary Figure 3 shows example output traces for the right ICA-MCA sections of all four models, respectively. As seen, conversion to super-resolution data mitigates the underestimation bias otherwise observed in the low resolution input data.

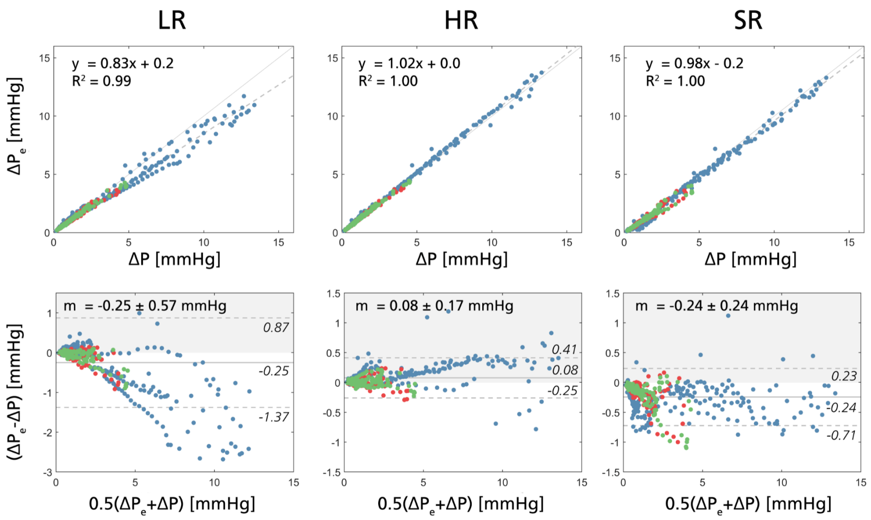


**Supplementary Figure 2** – Linear regression (top row) and Bland-Altman plots (bottom row), comparing relative pressure estimates to reference CFD equivalents using low resolution data (LR, 1 mm, left column), high resolution data (HR, 0.5 mm, middle column), and super-resolution data (SR, converting 1 mm to 0.5 mm, right column). The colors depict different model sets (training in blue (Subject 1 and 2), validation in red (Subject 3a), testing in green (Subject 3b)).

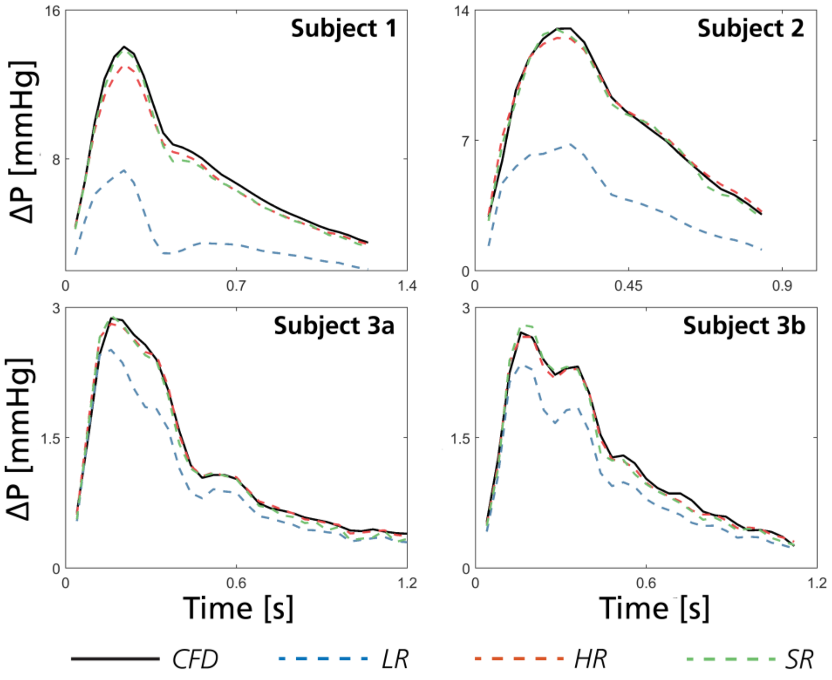


**Supplementary Figure 3** – Estimated relative pressures through the right ICA-MCA section in all subjects (left to right showing Subject 1, Subject 2, Subject 3a, and Subject 3b). In each graph, relative pressure estimates are derived from low resolution data in blue (LR, 1 mm), high resolution data in red (HR, 0.5 mm), super-resolved data in green (SR, converting 1 mm to 0.5 mm). True estimates are given by voxelized equivalents of the CFD pressure field generated at the HR sampling in black.

### C. Original vs. re-trained 4DFlowNet performance

#### 1) Estimation of super-resolution velocity

With primary results provided in Section III.A. of the main manuscript, Supplementary Figure 4 (next page) presented linear regression plots and coupled Bland-Altman representations for super-resolved cerebrovascular velocities using hyperparameters extracted from the original aortic 4DFlowNet. As described briefly in Section III.A.3, more distinct deviations are reported from ground truth high-resolution data, evident in both vessel core and vessel wall regions. Over all velocity components, a linear regression of about k = 0.88 (vessel core) or k = 0.78 (vessel wall) is given, distinctly lower than its re-trained equivalent. The original aortic network is however not associated with any specific estimation bias when it comes to super-resolved velocities (mean bias shift = -0.03 ± 0.11 and 0.01 ± 0.18 m/s for vessel core and vessel wall regions, respectively). Again echoing primary results reported in the main manuscript, increasing deviation are also evident with regard to peak velocity in both vessel core (MAE = 0.11 ± 0.08 m/s, relative error = 16.9 ± 0.1 %, cosine similarity = 0.99 ± 0.08) and vessel wall regions (MAE = 0.15 ± 0.11 m/s, cosine similarity = 0.93 ± 0.11).

#### 2) Estimation of super-resolution relative pressure

Coupling to 1) above, Supplementary Figure 5 (next page) presents linear regression plots and coupled Bland-Altman representations for super-resolved cerebrovascular relative pressures using hyperparameters extracted from the original aortic 4DFlowNet. As described in Section III.A.3, more distinct deviations are again reported from ground truth high-resolution data (k = 0.87 for super-resolution data, compared to k = 0.98 for high-resolution reference data). Also, in contrast to the re-trained network (see Section III.A.2), the original aortic network is also associated with relative pressure estimation bias (mean bias shift = -0.41 ± 0.58 mmHg for super-resolved data). Also echoing the data in the main manuscript, peak relative pressure estimates are given at a relative error of 14.8 ± 11.9 %, and a MAE of 0.4 ± 0.5 mmHg.

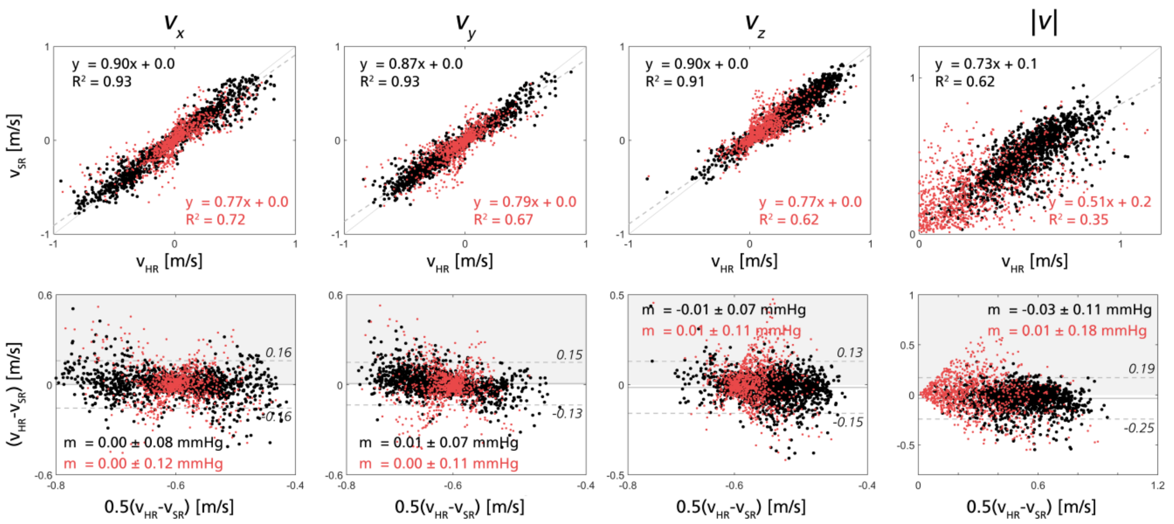


**Supplementary Figure 4 –** Top: Regression plot for each of the velocity components (Vx, Vy, and Vz) and velocity magnitude between ground truth and super-resolved image during the peak flow for in-silico test case (Subject 3b). Bottom: Bland-Altman plot for each of the velocity components during peak flow. The plots show 5% of the data points (randomly selected) within the vessel core (black) and vessel wall (red), respectively. All data generated using the original aortic 4DFlowNet.

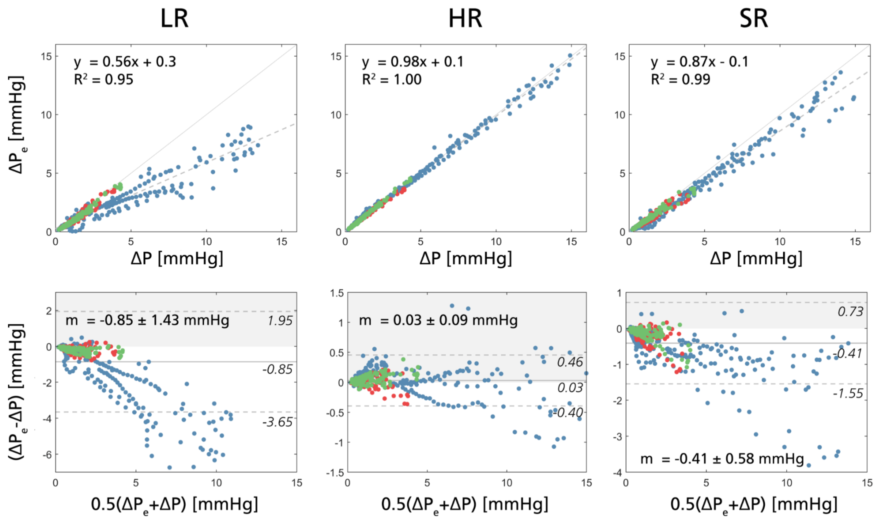


**Supplementary Figure 5 –** Linear regression (top row) and Bland-Altman plots (bottom row), comparing relative pressure estimates to reference CFD equivalents using low resolution data (LR, 1 mm, left column), high resolution data (HR, 0.5 mm, middle column), and super-resolution data (SR, converting 1 mm to 0.5 mm, right column) generated using the original aortic 4DFlowNet. The colors depict different data sets (training in blue (Subject 1 and 2), validation in red (Subject 3a), testing in green (Subject 3b)). Note that the LR and HR column are identical to Supplementary Figure 2.
